# AGAAT: Automated computational tool integrating different genotyping array and correctional methods for data analysis

**DOI:** 10.1101/2025.02.25.637414

**Authors:** Anil Prakash, Moinak Banerjee

**Author notes:** **Corresponding author:** Moinak Banerjee., Human Molecular Genetics Lab, Neurobiology and Genetics Division, Rajiv Gandhi Centre for Biotechnology, Thiruvananthapuram, Kerala, 695014, India.

## Abstract

Genotyping arrays are widely used in studying the genetics of complex diseases. Arrays with multi-ethnic markers enable population-scale and cost-effective genetic studies in underrepresented populations. But automated pipelines that convert raw data files into association results are lacking for these genotyping arrays. Also, combining variant files from multiple genotyping projects is difficult due to differences arising from multiple versions of standard file formats like Variant Calling Format (VCF). We have developed an automated tool named AGAAT (Automated Genotyping Array Analysis Tool) to overcome these problems. In addition to the automated pipelines for multi-ethnic arrays, we have added additional Python scripts for multiple testing correction using haplotype blocks, candidategene analysis, candidate-gene common-variant analysis and addition of 1000 genome phase-3 genotype counts to the controls, followed by case-control association analysis.

## INTRODUCTION

Large scale genetic association studies help in identifying significant risk variants of complex diseases with polygenic architecture. Genotyping arrays that identify common genetic variants are available for such studies with varying numbers of markers that are identified. Infinium Global Screening Array-24 (GSA) v3.0 is one such array with multi-ethnic and population specific markers covering the major global populations. Whereas, the Axiom Asia Precision Medicine Research Array (APMRA) targets multiple populations from East and South Asia. Raw data files generated by these genotyping arrays can be processed by Graphical User Interface (GUI) tools for converting them into vcf files. For large scale analysis, Command Line Interface (CLI) tools will be more practical [1, 2, 3]. But for genotypic array analysis, multiple tools have to be used while converting the raw data files into readable variant files, followed by association analysis.

A computational tool that automates everything from raw data conversion to association analysis will be helpful for large scale genotyping array analysis. Another problem that can arise in such situations is the availability of files from multiple genotyping projects. The raw data files can be analysed irrespective of such differences but standard files like vcf can vary due to their version differences. Merging separate datasets leads to errors, so incorporation of additional vcf files to already existing datasets can be challenging.

Another major difficulty in genome-wide association studies (GWAS) is multiple-testing correction. Bonferroni correction is commonly used in GWAS and it treats every genetic variant as an independent sample. This is not accepted in most cases because every genetic marker is not an independent unit due to differences in Linkage disequilibrium (LD) patterns.

In fact, multiple single nucleotide polymorphisms (SNP) in high LD can act as a single unit and this should be considered while doing multiple-testing correction.

Commercially available genotyping arrays will have variants in genes that are not important for the disease under study. Normal multi-marker association analysis will give equal importance to all the variants. This will increase the number of hypothesis tests for multiple-testing correction. Hence limiting the analysis to a set of pre-selected genes will improve the statistical power. Another way of improving the power is to increase the number of control samples. Control genotype data from large scale population genomic projects can be used for this purpose.

We have considered the above stated challenges in genotyping array analysis and developed our tool named AGAAT. It has automated bash scripts that automates raw data conversion, quality control, vcf file generation and case-control association. It can add additional vcf files to existing binary file sets. In addition to these pipelines, we have created Python scripts for LD based multiple-testing correction, candidate-gene and common-variant analysis. These scripts use the association and adjusted association output from PLINK for the extended analysis. We have incorporated Ensembl Rest-API into another script for adding genotype counts from 1000 genome phase-3 populations [4, 5].

## METHODS

The AGAAT tool contains bash scripts for conversion of raw .idat and .cel format files to vcf, followed by case-control association using PLINK. It can also add separate vcf files to the PLINK binary file set created from raw data files. The pipeline will check for errors in merging genetic variants and will exclude those variants before further association tests. Python scripts available with the tool will extract haplotype block output from PLINK and adjust the significance values based on the block and independent variants. The analysis can also be limited to a subset of genes which can be provided as user input. The list of candidate genes can be selected based on functional relevance and the disease under study. The analysis can also be limited to common variants within the input set of genes. Genotype counts from ethnically matching 1000 genome phase-3 populations [5] can be added to the control samples using AGAAT.

### GSA pipeline

gsa pipeline.sh combines all the tools [6, 7] that are desired for analyzing intensity data files to case-control association analysis as shown in Figure 1. Illumina Array Analysis Platform (IAAP) Genotyping CLI is used for converting raw .idat files to .gtc files [2]. The files thus generated are converted to a single vcf file using a bcftools plugin called +gtc2vcf [7]. The vcf file is used to create binary filesets for PLINK followed by case-control association analysis. The bash script combines all these tools in a sequential manner.

**Figure 1:**
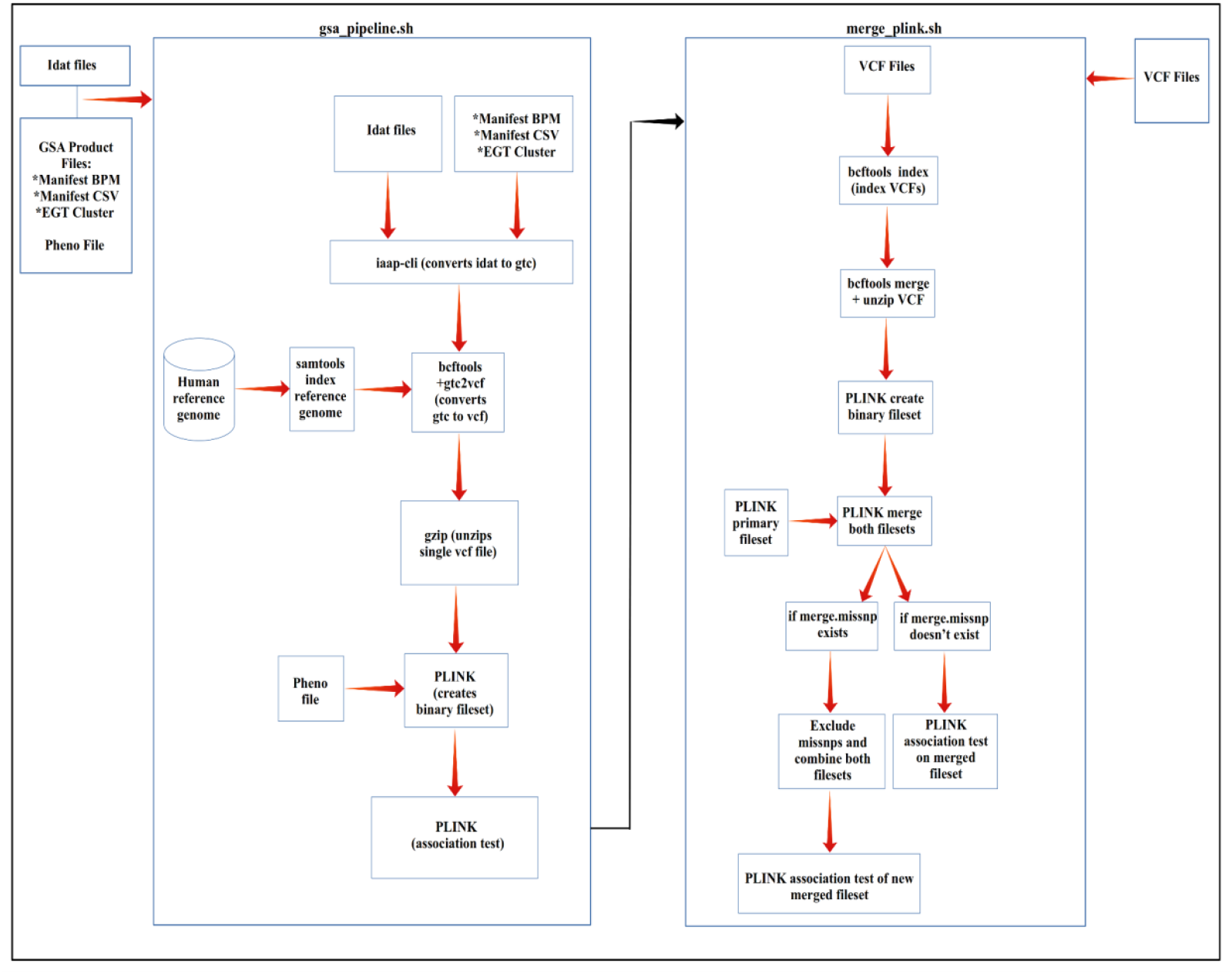
Flowchart diagram of AGAAT pipeline scripts.

The user needs to install the required tools and provide the input files accordingly. The directory having all the raw .idat files is required as input along with the reference genome and illumina product files. The product files specific to the array version are used as input. For case-control association analysis, an input text file with family and individual IDs of case samples are also required.

### Pipeline for merging separate datasets

The pipeline merges separate sets of vcf files to an existing PLINK binary fileset followed by case-control analysis. Combining vcf files from multiple genotyping projects can lead to errors due to differences in the versions and genetic variants. To overcome this, it is ideal to combine different datasets by a joint analysis of the raw data files. But in cases where raw files are absent this pipeline will help to add vcf file datasets to existing plink binary fileset. If error occurs due to merging of certain variants the script will exclude them and carry out association with the combined dataset (Figure 1).

### APMRA pipeline

The array provides genotyping output in the *.cel format and Analysis Power Tools (APT), which is a command line tool, provides separate functions for different analysis steps [3]. The quality control (QC) analysis functions will output values for different samples based on their quality. After these steps, the low-quality samples must be removed before further processing. Our pipeline automates all the APT functions that are required for APMRA analysis and filters samples according to the QC results (Figure 2).

**Figure 2:**
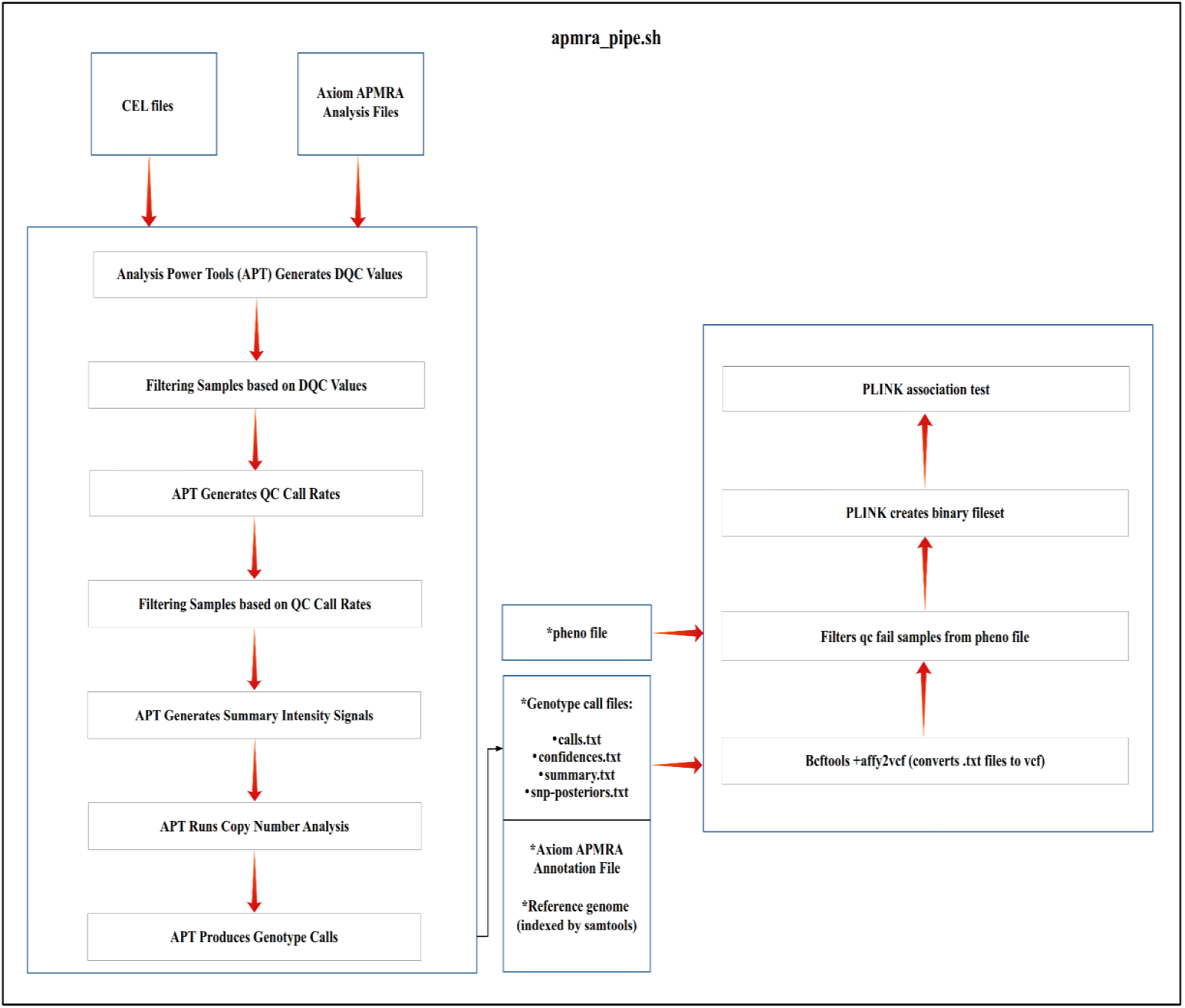
Flowchart diagram of APMRA pipeline.

The genotype calls in .txt formats from APT are converted to vcf using +affy2vcf bcftools plugin followed by case-control analysis using PLINK. The analysis and annotation files of the respective Axiom array must be given along with the input files. Additional vcf files can be added to the plink binary fileset using “merge plink.sh” bash script. The output association files generated by the pipeline can be given as input to the Python scripts for further processing.

### Candidate-gene and common-variant analysis

Candidate gene analysis helps in limiting the analysis to a subset of genes given as input. Normal genetic association will give equal importance to all the variants under study. Candidate gene analysis will limit the number of hypothesis tests and increase the statistical power required for detecting significant variants. Along with the input gene list, “common all 20180418.vcf” which is a dbSNP database of common variants are supplied as input [8]. All the common markers will be assigned to their respective genes by matching with this dataset.

The script will accept “plink.assoc.adjusted”, which is the adjusted association output file from PLINK and outputs gene-list based associations with Bonferroni correction (Figure 3).

**Figure 3:**
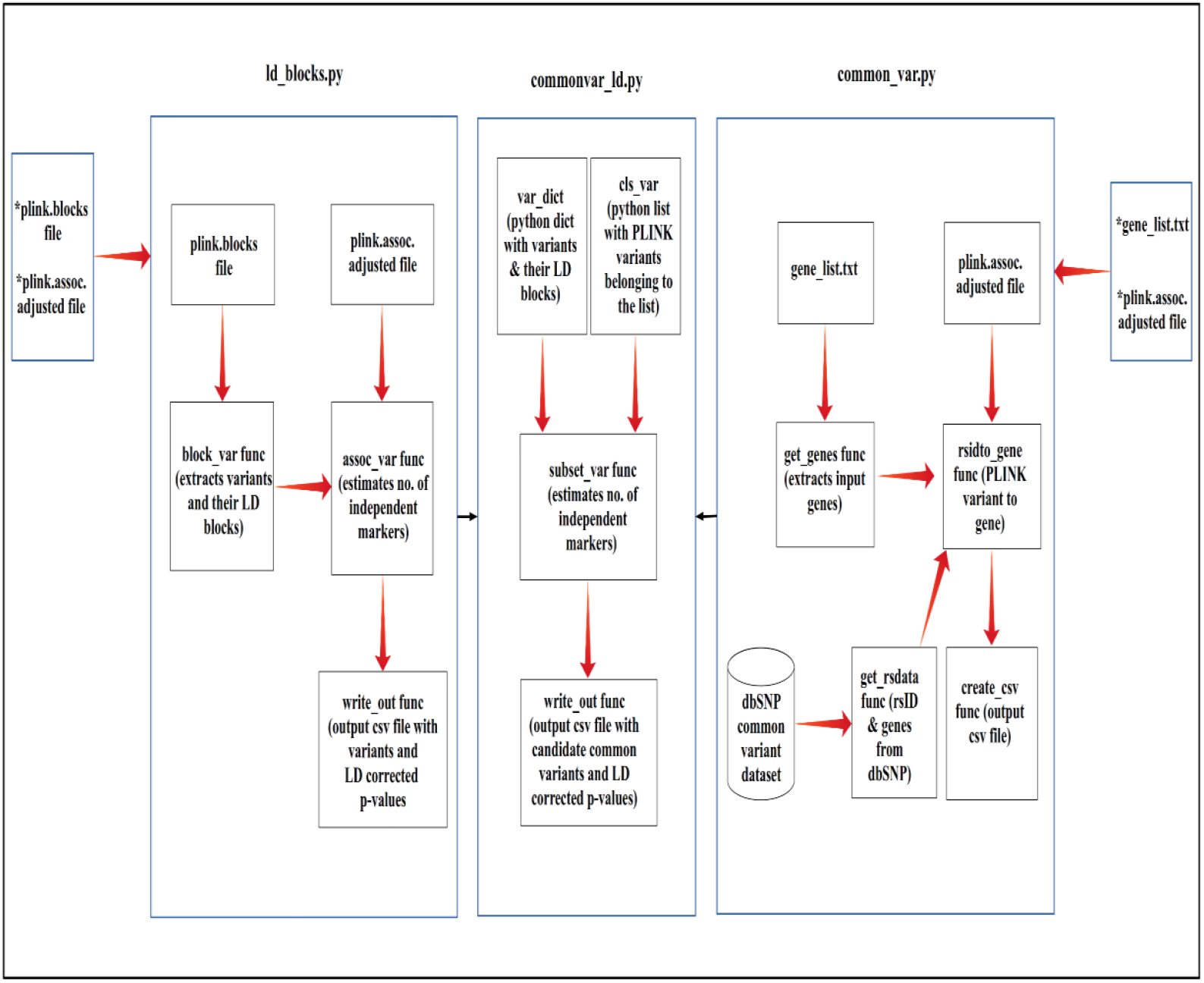
Flowchart diagram of python scripts for LD based multiple-testing correction and candidate-gene common-variant analysis.

### Multiple testing correction using LD blocks

GWAS analysis considers each marker as an independent sample, but in reality they are not, because of high LD between variants. The binary fileset given as input to PLINK can be used to generate haplotype blocks having markers within them. This script will consider haplotype blocks and markers outside them as independent units and carry out Bonferroni correction based on them. The script needs a “plink.blocks” file having haplotype blocks and variant IDs along with the adjusted association output file from PLINK (Figure 3). The output will have LD-block based adjusted associations.

### Candidate-gene common-variant analysis with LD based correction

The script combines LD-based multiple testing correction with candidate-gene common-variant analysis (Figure 3). LD based correction is done using haplotype blocks estimated using PLINK and input gene list along with dbSNP database, “common all 20180418.vcf”, are used for filtering the input variants.

### Candidate gene analysis

Filtering variants based on the dbSNP common variant database will remove rare variants from the association output. To avoid this, we have added another script named “candidate_gene.py” which annotates variants to their respective genes based on their coordinates. The bed file with genomic coordinates for all the genes can be extracted from the UCSC table browser [9]. The script will then select only those variants present in the candidate genes (Figure 4).

**Figure 4:**
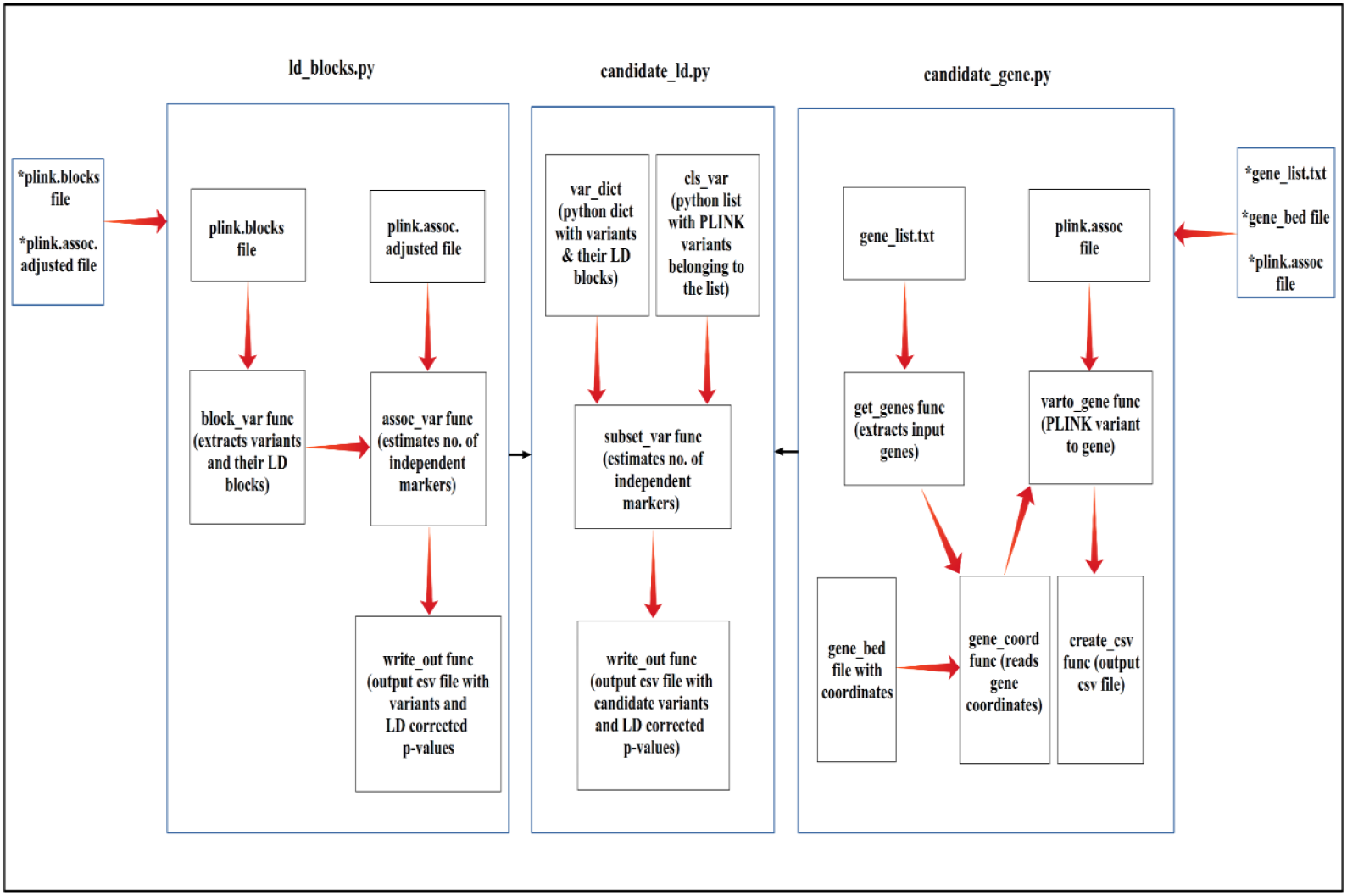
Flowchart diagram of python script for LD based multiple-testing correction and candidate-gene analysis.

### Combining LD based correction with candidate gene analysis

Python functions for multiple-testing correction using haplotype blocks derived from PLINK are available in “ld_blocks.py”. They were combined with functions defined in “candidate_gene.py” so that both candidate gene filtering and LD based correction were combined into a single script (Figure 4).

### Control genotype counts from 1000 Genome phase-3

The Control genotype count script adds genotype count data from 1000 genome phase-3 populations to the control sample counts [5]. This is followed by a repeated Hardy-Weinberg Equilibrium (HWE) test and case-control association. The 1000 genome populations and the sample population should be ethnically matching. The script uses Ensembl Rest API for extracting the genotype counts. The variants with added genotypes are frequently saved to file so that data retrieval can be resumed in case of errors caused by network connectivity issues. “plink.frqx”, which is the genotype count report file and “plink.assoc”, which is the association output file and the codes for 1000 genome phase-3 ethnically matched populations are given as input to the script (Figure 5).

**Figure 5:**
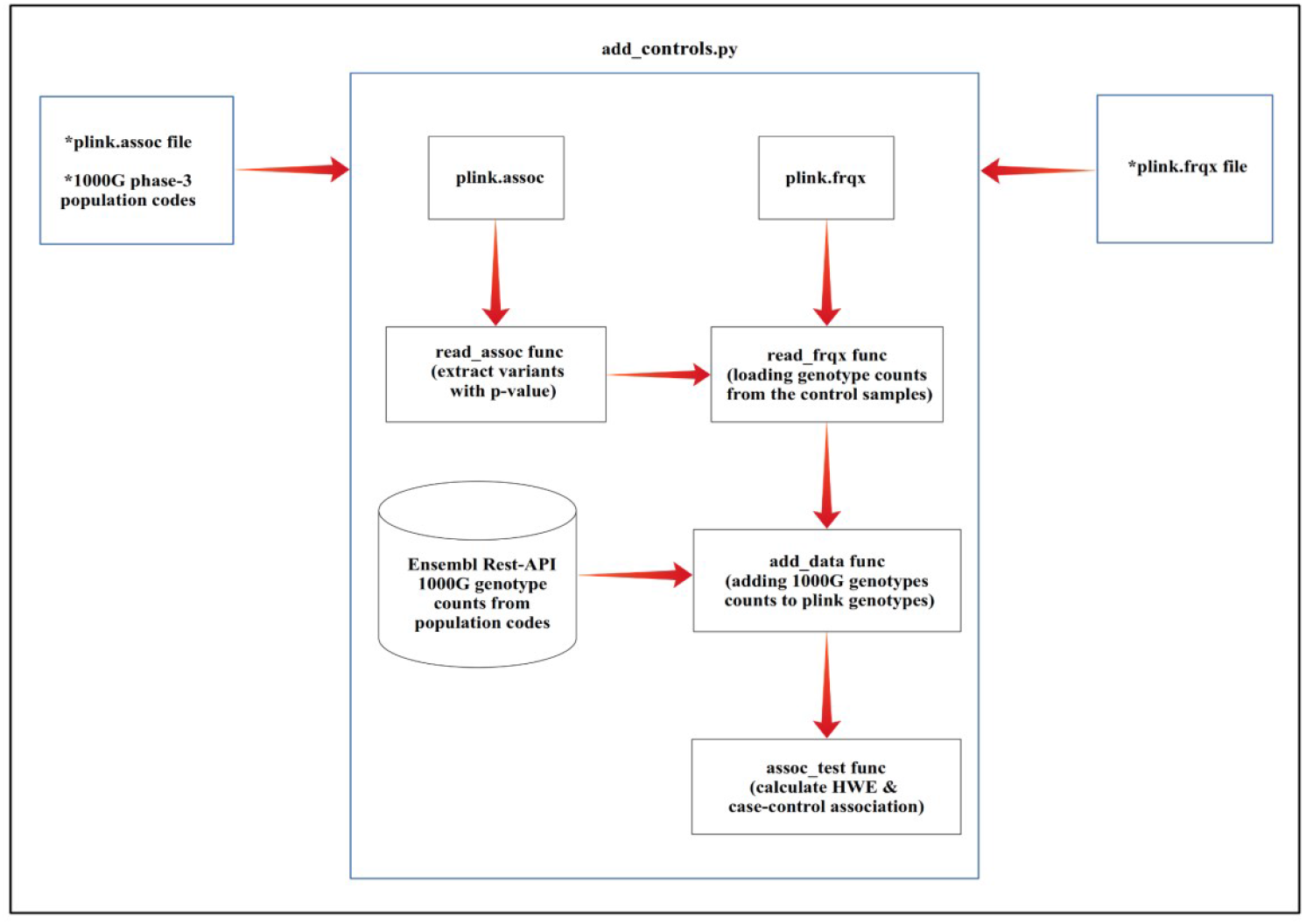
Flowchart diagram of python script for adding 1000 genome phase-3 genotype counts to input variants.

Adding the genotype counts can lead to conflicts due to different naming of alleles in Ensembl and PLINK output. In case of SNPs, Ensembl and PLINK major/minor alleles are matched against each other and for indels, they are matched based on the length of the allele names and frequency. It is advisable to manually re-check the genotype counts of the significant variants. Output text file from this script can be directly given as input to other python scripts used for purposes such as LD-based correction and candidate gene analysis.

Genotype counts of individual variants were retrieved from 1000 genome phase-3 populations using Ensembl Rest-API. For each input variant, a separate request to the Ensembl web server is needed. The total time taken will depend on the internet connectivity and the number of input variants. To reduce the time requirements, the input file, “plink.assoc”, can be sorted according to the raw p-value and the top list of variants can be saved into another text file and used as input.

## RESULTS AND DISCUSSION

The pipelines for converting raw output files of Illumina Infinium GSA v3.0 and Axiom APMRA genotyping arrays to association results and addition of another set of genotype data from vcf files are provided as bash scripts. They can be run in linux systems after installing the prerequisite tools. The input raw files and the product files for the respective arrays can be given as arguments to the scripts. Text file having information about the case samples should also be given as an argument for the association test. The rest of the scripts are written in Python and tested in Python version 3.8.8. The output files from the pipelines having association data can be used as input for the Python scripts. For annotating the variants to their respective genes, the common variant database from dbSNP and gene coordinates from UCSC table browser are used. Files generated by various PLINK commands are used as inputs for different scripts. For LD-based multiple testing correction, the haplotype blocks file generated by PLINK was used. The script, which adds genotype counts from 1000 genome populations, processes the control genotype count file generated by PLINK and compares the sample and 1000 genome genotype counts followed by HWE and association tests. Therefore, AGAAT is an integrated tool in the sense that it combines multiple analysis tools in the pipelines and the output thus generated serves as input for all the Python scripts.

The case-control association analysis of Autism Spectrum Disorder (ASD) in the Malayalam speaking Dravidian population was carried out using AGAAT. The study design involved Clinical diagnosis (DSM IV), Symptom evaluation and Demographic Evaluation for reviewing the samples. The study was limited by the sample size which included 156 cases and 111 controls. Control samples included ethnicity and sex matched individuals who had no family history of any neurological disorder. Genotyping was carried out by Illumina infinium GSA-24 v3.0 genotyping array. Raw data contained approximately 700,000 genetic markers for each sample. The Raw idat files from 156 cases and 111 controls were analysed using AGAAT which involved conversion into ped and map files using Illumina Array Analysis Platform (IAAP) Genotyping Command Line Interface (CLI) and association test using plink-1.90. The unadjusted p-values were corrected for multiple testing using Bonferroni adjustment (Supplementary Figure S1). A candidate gene based analysis was carried out by restricting the analysis to markers belonging to a candidate list of genes. This will reduce the multiple testing burden and improve the statistical power of the study.

The network analysis was limited to a list of Circadian genes [10] (Supplementary Figure S2) and the three significant SNPs that were detected after multiple testing corrections (rs2859388, rs228642 and rs707467) (Supplementary Figure S1). They belong to the PER3 gene in chromosome number 1 and both rs2859388 and rs228642 are reported to be associated with Bipolar affective disorder as a multi-marker haplotype [11]. The rs228642 polymorphism is found to be interacting with stressful life events to influence sleep patterns in females [12].

AGAAT is a bioinformatic tool with automated pipelines and python scripts that can be used for extended analysis using the association results. Use of GUI tools will be time-consuming for large scale genotyping projects. AGAAT makes it easier to work with a huge number of raw data files and integrates different tools in a sequential manner.

The adjusted association output file from PLINK is used as an input for the python scripts. This significantly reduces the time and effort because it eliminates the requirement of repeating association tests. The scripts goes through the variants listed in the association output and assigns gene names to them using publicly available dbSNP common variant databases or using a bed file with gene coordinates. The haplotype blocks that are used for LD based correction are generated from PLINK using the input dataset itself. This will eliminate the discrepancies in LD between variants that can arise due to population differences.

AGAAT is a tool with robust pipelines and scripts for genotyping array analysis. It is open source and available on GitHub. Researchers can run the bash script pipelines in Linux systems for analysing bulk genotyping array datasets. The tool is available for developers to modify as per the emerging needs and future developments. Overall, AGAAT will be a useful tool for multi-ethnic and population specific genotyping array analysis. Candidate-gene and common-variant analysis will be helpful for genetic studies concentrating on a set of functional genes and those with limited sample size. It will aid in the study of genetics and complex diseases in neglected populations with fewer available samples.

## DATA AVAILABILITY

AGAAT is freely available at https://github.com/anilprakash94/agaat/.

## ACKNOWLEDGEMENTS

We are grateful to the Dept. of Biotechnology, Govt. of India and Council of Scientific and Industrial Research (CSIR) for Senior Research Fellowship (to A.P.).

## AUTHOR’S CONTRIBUTIONS

A.P. and M.B. conceptualized the work, developed the tool and wrote the manuscript.

---

**Figure S1:**
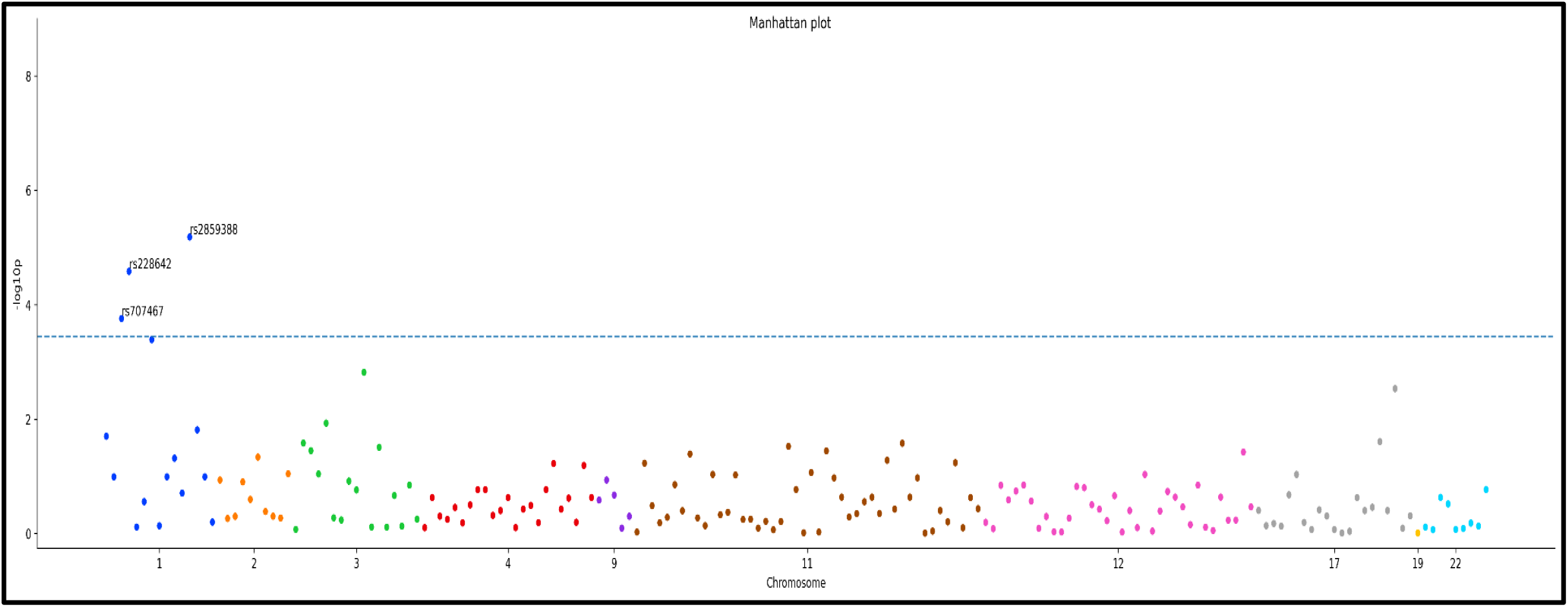
Candidate gene association results. Scatter plot of circadian genes which were selected as candidates.

**Figure S2:**
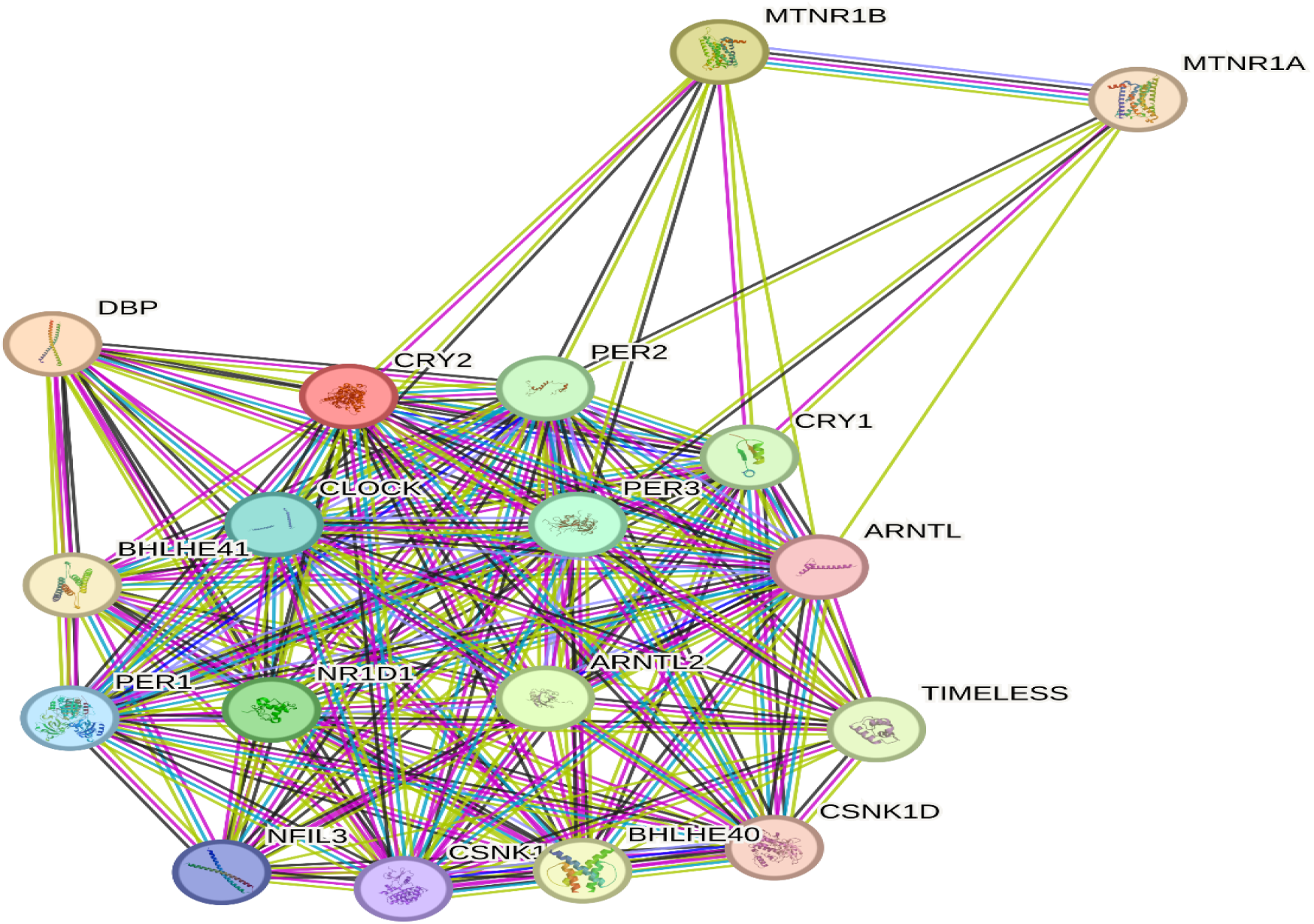
STRING Protein-Protein Interaction Network of candidate genes. Network diagram of circadian genes which were selected as candidates.

## REFERENCES

1. Chang CC, Chow CC, Tellier LC, Vattikuti S, Purcell SM, Lee JJ. Second-generation PLINK: rising to the challenge of larger and richer datasets. Gigascience. 2015;4:7. doi: 10.1186/s13742-015-0047-8.

2. IAAP Genotyping CLI v1.1. Illumina Array Analysis Platform Genotyping Command Line Interface v1.1. [Software]. https://sapac.support.illumina.com/downloads/iaap-genotyping-cli.html

3. Affymetrix, Inc. [Software]. https://www.thermofisher.com/

4. Yates A, Beal K, Keenan S, McLaren W, Pignatelli M, Ritchie GR, Ruffier M, Taylor K, Vullo A, Flicek P. The Ensembl REST API: Ensembl Data for Any Language. Bioinformatics. 2015;31(1):143–5. doi: 10.1093/bioinformatics/btu613.

5. 1000 Genomes Project Consortium; Auton A, Brooks LD, Durbin RM, Garrison EP, Kang HM, Korbel JO, Marchini JL, McCarthy S, McVean GA, Abecasis GR. A global reference for human genetic variation. Nature. 2015;526(7571):68–74. doi:10.1038/nature15393.

6. Danecek P, Bonfield JK, Liddle J, Marshall J, Ohan V, Pollard MO, Whitwham A, Keane T, McCarthy SA, Davies RM, Li H. Twelve years of SAMtools and BCFtools. Gigascience. 2021;10(2):giab008. doi: 10.1093/gigascience/giab008.

7. gtc2vcf. [Software]. https://github.com/freeseek/gtc2vcf.

8. Sherry ST, Ward MH, Kholodov M, Baker J, Phan L, Smigielski EM, Sirotkin K. dbSNP: the NCBI database of genetic variation. Nucleic Acids Res. 2001;29(1):308–11. doi:10.1093/nar/29.1.308.

9. Karolchik D, Hinrichs AS, Furey TS, Roskin KM, Sugnet CW, Haussler D, Kent WJ. The UCSC Table Browser data retrieval tool. Nucleic Acids Res. 2004;32(Database issue):D493–6. doi:10.1093/nar/gkh103.

10. Szklarczyk D, Gable AL, Lyon D, et al. STRING v11: protein-protein association networks with increased coverage, supporting functional discovery in genome-wide experimental datasets. Nucleic Acids Res. 2019 Jan 8;47(D1):D607–D613.

11. Nievergelt CM, Kripke DF, Barrett TB, et al. Suggestive evidence for association of the circadian genes PERIOD3 and ARNTL with bipolar disorder. Am J Med Genet B Neuropsychiatr Genet. 2006 Apr 5;141B(3):234–41.

12. Antypa N, Mandelli L, Nearchou FA, et al. The 3111T/C polymorphism interacts with stressful life events to influence patterns of sleep in females. Chronobiol Int. 2012 Aug;29(7):891–7.

